# The potential aroma and flavor compounds in *Vitis* sp. cv. Koshu and *V. vinifera* L. cv. Chardonnay under different environmental conditions

**DOI:** 10.1101/383000

**Authors:** Sharon Marie Bahena-Garrido, Tomoko Ohama, Yuka Suehiro, Yuko Hata, Atsuko Isogai, Kazuhiro Iwashita, Nami Goto-Yamamoto, Kazuya Koyama

## Abstract

**ABSTRACT:** *BACKGROUND:* Koshu, a hybrid of *Vitis vinifera* L. and *V. davidii* Foex, is the most popular indigenous cultivar for wine production in Japan. However, little is known about the potential aroma compounds it contains and how environmental factors affect these. In this study, we obtained comprehensive profiles of the volatile (both glycosidically bound and free) and phenolic compounds that occur in Koshu berries, and compared these with similar profiles for *V. vinifera* cv. Chardonnay. We then compared the response of these two cultivars to bunch shading and the ripening-related phytohormone abscisic acid (ABA).

*RESULTS:* Koshu berries contained significantly higher concentrations of phenolic compounds, hydroxycinnamic acid derivatives, and some volatile phenols, such as 4-vinyl guaiacol and eugenol, than Chardonnay berries, which are thought to contribute to the characteristics of Koshu wine. In addition, Koshu berries had a distinctly different terpenoid composition from Chardonnay berries. Shading reduced the concentrations of norisoprenoid in both cultivars, as well as several phenolic compounds particularly their volatile derivatives in Koshu. The exogenous application of ABA induced ripening and increased the concentrations of lipid derivatives, such ashexanol, octanol, nonanol, and 1-octen-3-ol. Linear discriminant analysis showed that the aromatic potential could be discriminated clearly based on cultivar, bunch shading, and ABA application.

*CONCLUSION:* The unique secondary metabolite profiles of Koshu and their different responses to environmental factors could be valuable for developing various styles of Koshu wines and new cultivars with improved quality and cultural characteristics.

## INTRODUCTION

In wine production, aroma is an important component of grape quality and wine flavor. Several volatile organic compounds that affect flavor and aroma have been identified in grapes^1^, many of which are synthesized at the end of the ripening stage during berry development^2,3^. Many of the aroma compounds in the berries are produced as volatile, odor-active forms (or free volatiles) and non-volatile, conjugated forms (or bound volatiles), which are cysteinylated, glutathionylated, or most commonly glycosylated precursors. Aglycones are broadly classified as terpenoids, C_13_-norisoprenoids, thiols, shikimates, and so on^1,4–8^. All of these non-volatile precursors serve as a potential aroma reservoir and are converted into various flavors by the yeast enzymes during fermentation, as well as by acidic fermentation and aging conditions^9,10^. They are released from their conjugated forms through hydrolysis and eventually develop into aroma compounds in wine. Thus, a complete understanding of grape aromas in wine requires knowledge of the distribution of both free and glycosylated volatiles, as well as the composition of the various odorants that are derived from glycoside hydrolysis^8,11^.

The accumulation of flavor compounds and volatile aromas is affected by several environmental factors, such as soil type, water availability, sunlight, and temperature^12–15^. Among these factors, sunlight appears to have the greatest impact, with several studies having shown that the synthesis of aroma compounds such as monoterpenoids and C_13_ norisoprenoids, as well as anthocyanins and tannins, in red wine grapes is promoted by manipulating light exposure around the bunches^13,16,17^. Increases in the overall concentrations of phenolic compounds are considered to have negative effects on the flavor properties of some white wine cultivars. However, comprehensive analyses of the secondary metabolite contents, including phenolic and aroma compounds in both free and bound forms, will allow us to conduct a holistic evaluation of the impact of light exposure on the quality of wine grapes.

Abscisic acid (ABA) is known to be both a stress-and ripening-related hormone in grapevines^18^, with exogenous ABA application not only inducing the accumulation of anthocyanin but also affecting various ripening-related metabolic processes in the skins of grapes^19^. In addition, it has recently been reported that ABA application increases the concentration of C_6_ aroma compounds derived from fatty acids and mono- and sesquiterpenes^20,21^. However, it is not yet fully understood how ABA affects secondary metabolites other than flavonoids, such as volatile compounds, particularly in white wine cultivars.

In Japan, grape and wine production is becoming increasingly popular and important. Among the grape cultivars that are used for winemaking, Koshu is one of the most popular, with a total of 4,568 tons (representing 17.3% of production) having been produced in 2015, mainly from Yamanashi Prefecture, compared with 1,243 tons (5%) for Chardonnay, which is mostly grown in Yamanashi and Nagano Prefectures (Source: Japanese National Tax Agency). Koshu is an indigenous, purple-skinned grape that has been cultivated in Central Japan for more than 1,000 years for use as table grapes and more recently for white winemaking. Goto-Yamamoto *et al*.^22^ recently reported that Koshu is a hybrid of *Vitis vinifera* L. (70%) and the Chinese wild species *V. davidii* Foex or closely related species (30%) based on single nucleotide polymorphism analyses and partial sequencing of the chloroplast DNA.

Koshu wine has traditionally been considered to be neutral and relatively indistinctive or “moderate.” However, it has recently been found that ß-damascenone and 3-mercaptohexan-1-ol (3MH) contribute to its aroma, leading to the development of aromatic Koshu wines with enhanced fruity or citrus characters through viticultural practices focusing on these flavor precursors in the berries and enological techniques^23,24^. However, more detailed elucidation of the potential flavor compounds in this cultivar and an understanding of how these compounds are modulated by ripening stage and environmental cues will allow its full potential to be realized.

In this study, we performed comprehensive analyses of the secondary metabolites in Koshu berries, including aroma (or volatiles) compounds in both bound and free forms and phenolic compounds, and compared these with the compounds that occur in the pure *V. vinifera* cultivar Chardonnay, which is also non-aromatic and used to produce white wine. We then compared the response of these two cultivars to bunch shading and ABA application. To the best of our knowledge, this is the first attempt to characterize the enological potential of Koshu by focusing on its flavor precursors, and the findings of this study will provide us with new insights into approaches required for improving the quality of wine produced by this non-aromatic hybrid of Eastern and Western cultivars.

## MATERIALS AND METHODS

### Chemicals and reagents

Aroma and flavor standards were purchased from Tokyo Chemical Industry Co., Ltd. (TCI), Japan, and Wako Co., Ltd (WAKO), Japan. Dichloromethane, liquid chromatography–mass spectrometry (LCMS)-grade methanol, citric acid, phosphates, and NaOH were obtained from WAKO. Pure water was obtained from a Milli-Q purification system (Millipore, Bedford, MA, United States).

### Plant and fruit materials, field experiments, and sampling

This study was conducted in 2013 using 20-year-old Koshu and Chardonnay grapevines that had been cultivated under the same management conditions (*i.e*., training, pruning, irrigation, soil, and fertility) in the experimental vineyard at the National Research Institute of Brewing, Higashi-Hiroshima City, Hiroshima Prefecture, Japan. Each of the vines had been trained on a Cordon trellising system^25^ and bore 15–20 bunches of berries. The vines of each cultivar were divided into three plots to provide three biological replicates and the bunches of each plot were subjected to treatments.

A shading treatment was applied to berry bunches of both cultivars 2 weeks before véraison using a light-proof box that was identical to that used by Koyama *et al*.^26^ This was designed to completely eliminate light through the use of black polypropylene sheeting (0.7 mm) while maximizing airflow and so was considered adequate for examining the impact of light exclusion without affecting the temperature. In 2013, véraison began around August 15 for Koshu and July 16 for Chardonnay.

During the véraison stage, some of the bunches were exogenously treated with 400 mg/L abscisic acid (S-ABA) solution (Sigma-Aldrich, Inc., St. Louis, MO, United States) by spraying, following the method of Mori *et al*.^27^

At 3 and 6 weeks after véraison (WAV), 200 fresh berries were randomly sampled from each treated bunch per plot, taking into account the number of berries per cluster and ensuring that the number of berries taken from the shaded and sun-exposed sides was equal. The ripeness indices were then analyzed following Koyama *et al*,^28^ while other fractions of the samples were deseeded, immediately frozen in liquid nitrogen, and stored at −80°C until further analysis.

### Analysis of bound and free volatile organic compounds

#### Berry sample preparation

Volatile compounds were extracted from 100 whole berries from each of the three replicate groups by partially crushing the berries in liquid nitrogen using a pestle and mortar and then powdering them in a bead miller (Yasui Kikai, Japan). Approximately 10 g of the powdered berries was then used for extraction following the method of Loscos *et al*.^29^ with various modifications. Extraction was carried out by adding 0.13 M NaF, 50 mg/L of S-ABA solution, and 50 ng/L of the internal standard n-heptyl β-D-glucopyranoside (Sigma-Aldrich) to each berry sample and shaking for 15 min at 40°C. The mixture was then centrifuged and filtered through a 5-μm membrane filter (Sartorius, Goettingen, Germany).

#### Extraction of the bound and free volatiles

Volatiles were extracted by using a tandem extraction method, a modified version of the method of Loscos *et al*.^29^ and Ibarz *et al*.^30^ which mainly used the solid-phase extraction combined with stir bar sorptive extraction (SBSE) method. SBSE, which uses a stir bar coated with polydimethylsiloxane (PDMS), has been widely used to extract volatiles due to its versatility and sensitivity^31^. Briefly, the berry extracts were applied to LiChrolut^®^-EN solid-phase extraction cartridges (Merck, Germany) that had been pre-conditioned according to the manufacturer’s protocol. The bound volatiles were eluted with an ethyl acetate:methanol (9:1, v/v) mixture and the eluate was divided into two for enzymatic hydrolysis and heat-acid hydrolysis and then concentrated to dryness. Enzymatic hydrolysis was used to characterize the pool of bound fractions, as this does not induce any further transformation in the chemical structure of the aglycones that are released. However, like previous studies,^32,3^ we found that this method could not detect norisoprenoids, which are thought to contribute significantly to the aroma of non-aromatic cultivars.^11,34^ Therefore, we analyzed the bound form of C13-norisoprenoids separately using heat-acid hydrolysis by adding 0.2 M citric acid buffer solution at pH 2.5 to the sample and heating in a water bath to 100°C for 1 h in an encapsulated vial under a nitrogen atmosphere.^29,30^ Any released volatiles were then further concentrated by SBSE following the protocol of Isogai *et al*.^31^ with some modification. Briefly, the eluent was dispensed into a 10-mL glass headspace vial containing 2 g of NaCl and the internal standard 3-octanol (0.5 mg/L) with a Twister^®^ (Gerstel, Mulhein an der Ruhr, Germany; 10-mm long, 0.5-mm layer of PDMS). For the analysis of free volatiles, the berry extracts were used directly for SBSE. All the extracts were further analyzed by gas chromatography–mass spectrometry (GC-MS).

#### GC–MS analysis

Each Twister was placed in a desorption tube, which was introduced into a thermal desorption system by a TDSA2 auto-sampler (Gerstel). The stir bar was then thermally desorbed by heating the TDS from 20°C (1 min) to 230°C (for 4 min) at a rate of 60°C/min. The desorbed components were cryofocused in the CIS4 at −150°C, following which the CIS4 temperature was increased to 250°C at a rate of 12°C/s and held for 10 min. The trapped components were then injected into a GC column using the splitless mode.

GC–MS was performed using an Agilent 6890N/5973 GC–MS system equipped with an HP-INNOWax column (30 m × 0.25 mm × 0.25 μm film thickness; Agilent Technologies Inc.). The GC oven temperature was programmed to start at 40°C with a 3-min hold and then to increase to 240°C at a rate of 5°C/min and to hold at 240°C for 10 min. Helium was used as a carrier gas at a constant flow rate of 1.0 mL/min. For the mass spectrometry (MS) system, the transfer line, quadrupole, and ionization source were set at 250, 150, and 230°C, respectively, and electron impact mass spectra were recorded at 70 eV ionization voltages. The acquisitions were performed in full-scan mode (35–400 *m/z*).

Peak area was normalized against the 3-octanol standard. Compound identifications were made by matching the mass spectra to the National Institute of Standards and Technology (NIST) 2011 library and retention indices that were calculated by analyzing C7–C27 n-alkane standards under the same chromatographic conditions. When authenticated standards were not available, tentative identifications were based on the NIST 2011 library and a comparison of retention indices reported in the literature. Selective ion-monitoring mass spectrometry was used to quantify the aroma compounds.

### Extraction and quantification of proanthocyanidins and monomeric phenolic compounds

Proanthocyanidins (PAs) and monomeric phenolic compounds were extracted simultaneously from the whole berries, excluding the seeds, following the modified method of Mané *et al*.^35^ Methods for the identification and quantification of each individual constitutive unit within PAs and phenolic monomers and the determination of their total concentrations, and details of the high-performance liquid chromatography conditions have been described previously.^28,36^ Each compound was expressed as the amount per gram of whole berry fresh weight (μg/g of whole berry FW).

### Statistical analyses

In all experiments, multi-way analysis of variance was used to analyze the effect of cultivar and treatment on the volatile and phenolic compounds and Tukey’s honest significant difference (*p* < 0.05) test was used to further investigate any significant effects of treatment within each cultivar. In addition, least square means Student’s *t*-tests were used to examine whether there were significant differences in the concentration of a particular volatile component between cultivars in the control treatments at 3 and 6 WAVs. The distribution of the groups of the bound volatiles from each untreated cultivar at 6 WAV was compared by means of the Chi-square test, *p* < 0.05 to determine significant difference between cultivars.

Principal component analysis (PCA) was performed to investigate the relationships between the bound volatile constituents of the two different cultivars and the different treatment groups. Linear discriminant analysis (LDA) was also carried out to discriminate the flavor precursor compositions of the berry samples, whereby we established a single model for discriminating the different treatment groups using the selected volatile precursors by forward stepwise regression. The accuracy of classification was then verified using leave-one-out cross validation.^37^ All the statistical analyses were carried out using the JMP ver. 13.0 software (SAS Institute Inc., Cary, NC, United States).

## RESULTS

### Maturation of Koshu and Chardonnay berries

#### Cultivar differences

Koshu berries were approximately twice the weight of Chardonnay berries under the growing conditions used in this study (Table 1). In addition, Koshu berries had lower total soluble solid (TSS) concentrations (°Brix) and higher total acidity (TA) levels than Chardonnay berries, which are innate characteristics of this cultivar. Sampling was carried out at around 6 WAV, when the increase in TSS had plateaued in the control treatments and both cultivars were considered to have reached full maturity.

**Table 1.**
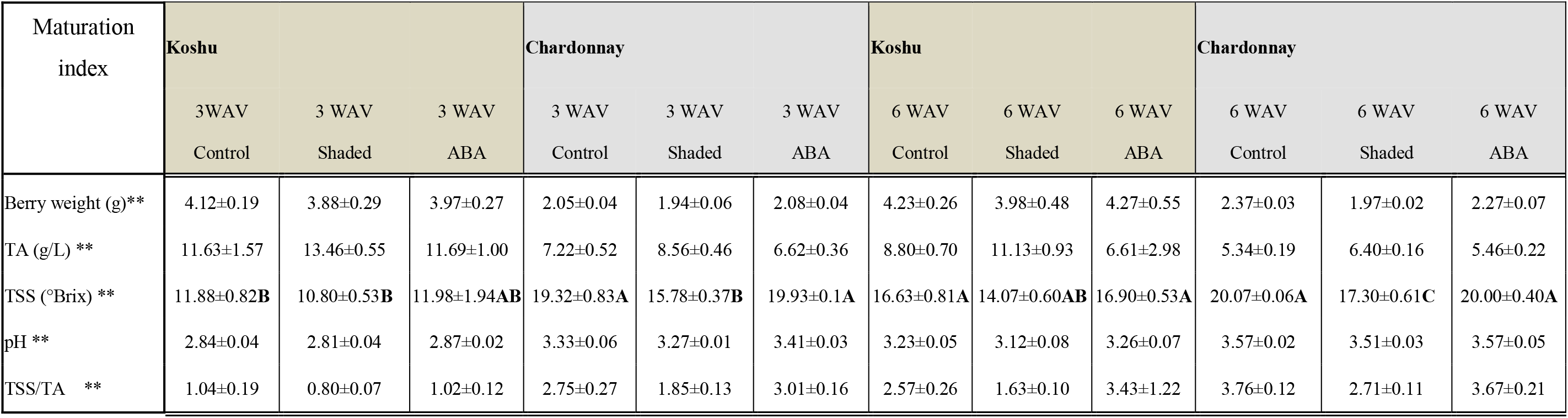
Maturity of Koshu and Chardonnay berries among treatments as measured using the maturation indices berry weight, total acidity (TA), total soluble solid (TSS) concentrations (Brix), TSS/TA ratio, and pH. Berries were measured at 3 and 6 weeks after véraison (WAV) and were treated with bunch shading, the exogenous application of abscisic acid (ABA), or no treatment (control). ** Maturation indices of Koshu and Chardonnay were significantly different (least square means Student’s t-test, *p* < *0.01*). Values are means ± standard errors. Different letters indicate significant differences among treatments within each cultivar (Tukey’s honest significant difference test, *p* < *0.01*).

| Maturation index | Koshu |  |  | Chardonnay |  |  | Koshu |  |  | Chardonnay |  |  |
| --- | --- | --- | --- | --- | --- | --- | --- | --- | --- | --- | --- | --- |
|  | 3WAV | 3 WAV | 3 WAV | 3 WAV | 3 WAV | 3 WAV | 6 WAV | 6 WAV | 6 WAV | 6 WAV | 6 WAV | 6 WAV |
|  | Control | Shaded | ABA | Control | Shaded | ABA | Control | Shaded | ABA | Control | Shaded | ABA |
| Berry weight (g)** | 4.12±0.19 | 3.88±0.29 | 3.97±0.27 | 2.05±0.04 | 1.94±0.06 | 2.08±0.04 | 4.23±0.26 | 3.98±0.48 | 4.27±0.55 | 2.37±0.03 | 1.97±0.02 | 2.27±0.07 |
| TA (g/L) ** | 11.63±1.57 | 13.46±0.55 | 11.69±1.00 | 7.22±0.52 | 8.56±0.46 | 6.62±0.36 | 8.80±0.70 | 11.13±0.93 | 6.61±2.98 | 5.34±0.19 | 6.40±0.16 | 5.46±0.22 |
| TSS (°Brix) ** | 11.88±0.82**B** | 10.80±0.53**B** | 11.98±1.94**AB** | 19.32±0.83**A** | 15.78±0.37**B** | 19.93±0.1**A** | 16.63±0.81**A** | 14.07±0.60**AB** | 16.90±0.53**A** | 20.07±0.06**A** | 17.30±0.61**C** | 20.00±0.40**A** |
| pH ** | 2.84±0.04 | 2.81±0.04 | 2.87±0.02 | 3.33±0.06 | 3.27±0.01 | 3.41±0.03 | 3.23±0.05 | 3.12±0.08 | 3.26±0.07 | 3.57±0.02 | 3.51±0.03 | 3.57±0.05 |
| TSS/TA ** | 1.04±0.19 | 0.80±0.07 | 1.02±0.12 | 2.75±0.27 | 1.85±0.13 | 3.01±0.16 | 2.57±0.26 | 1.63±0.10 | 3.43±1.22 | 3.76±0.12 | 2.71±0.11 | 3.67±0.21 |

#### Effects of bunch shading and ABA

Shading resulted in lower TSS concentrations, higher TA levels, and lower TSS/TA ratios in both cultivars, but particularly in Koshu at 6 WAV and Chardonnay at 3 and 6 WAVs (Table 1). The exogenous application of ABA resulted in a stronger yellowish color in the skins of Chardonnay berries at 6 WAV, suggesting accelerated ripening. However, this had no significant effect on the other maturation indices (*i.e*., berry weight, TSS, TA, and pH) in either cultivar.

### Compositions of the bound volatiles in Koshu and Chardonnay berries

#### Cultivar differences

In total, we identified approximately 77 precursors of the volatile compounds (Table 2) and further categorized these according to their biological origin, which placed the volatiles into four large groups of components, *i.e*., lipid derivatives, benzene derivatives, terpenoids, and norisoprenoids.^30^ There were remarkable differences in the composition of bound volatiles between the two cultivars, with Koshu berries containing significantly higher ratios of volatile phenols and vanillins, and Chardonnay berries containing significantly higher ratios of norisoprenoids and benzoids (Figure 1). These differences in bound volatile compositions of untreated berries at harvest can be viewed in the heat map provided in Figure 2. The major terpenoids in Koshu berries were alpha-terpineol, linalool oxide pyranoside, and geraniol, while Chardonnay berries contained linalool, cis-linalooloxide, nerol, and geraniol (Table 2). A closer inspection of the results shows that, interestingly, Koshu berries also possessed a larger number of minor terpenoids, such as terpinen-1-ol, terpinen-4-ol, cis-carveol, pino-carveol, myrtenol, and linalool oxide pyranoside, which was distinctively different from Chardonnay berries, as well as higher concentrations of vitispirane A and ß-damascenone in norisoprenoids. By contrast, Chardonnay berries were found to have higher accumulation levels of total C_13_-norisoprenoids and significantly higher concentrations of the other bound norisoprenoids such as 1,1,6,-trimethyl-1,2-dihydronapthalene (TDN), 4-(2,3,6-trimethylphenyl)buta-1,3-diene (TPB), and 3-oxo-alpha-ionol at 3 WAV, and actinidols in the early and late ripening stages than Koshu berries (Figure 2).

**Figure 1.**
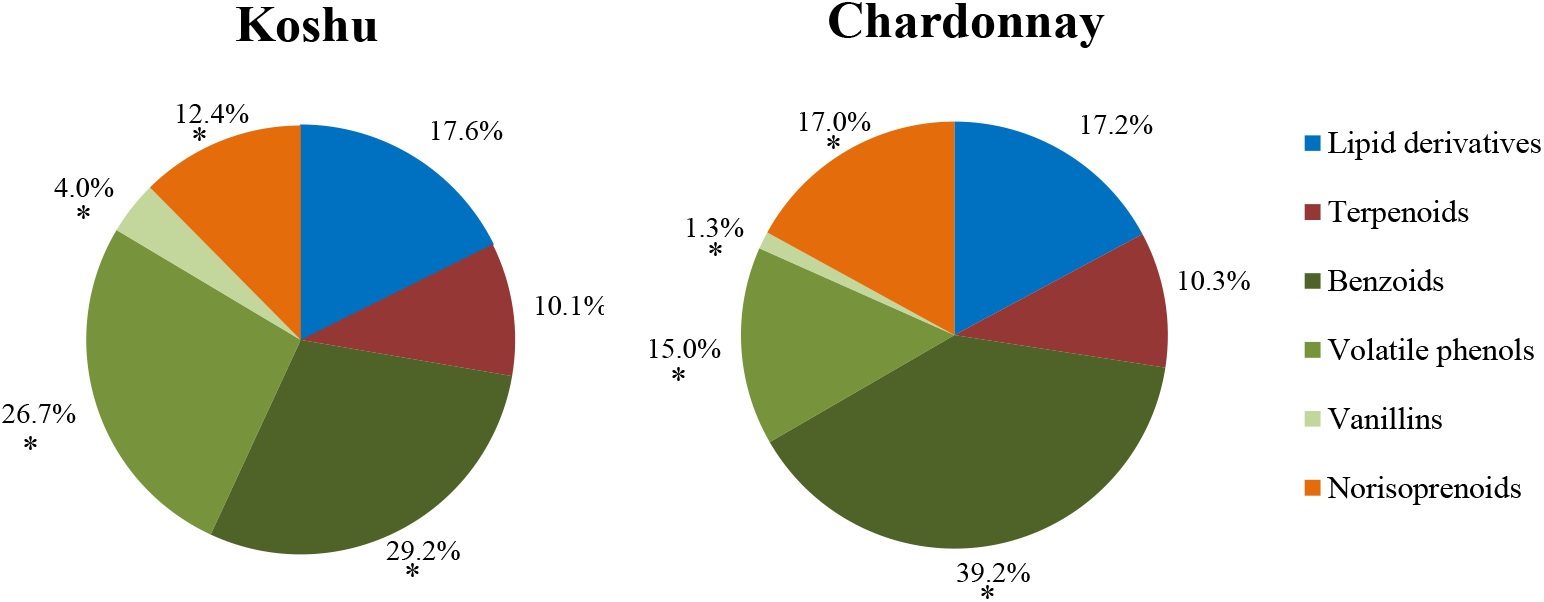
Composition of the bound volatiles in untreated Koshu and Chardonnay berries at the harvest stage (6 weeks after véraison [WAV]). The detected compounds were classified as lipid derivatives, terpenoids, benzoids, volatile phenols, vanillins, and norisoprenoids. The concentration of each group of compounds was determined by summing the amounts shown in Table 2. Data are means (n = 3). * Significant difference between groups (Chi-square test, *p* < 0.05).

**Figure 2.**
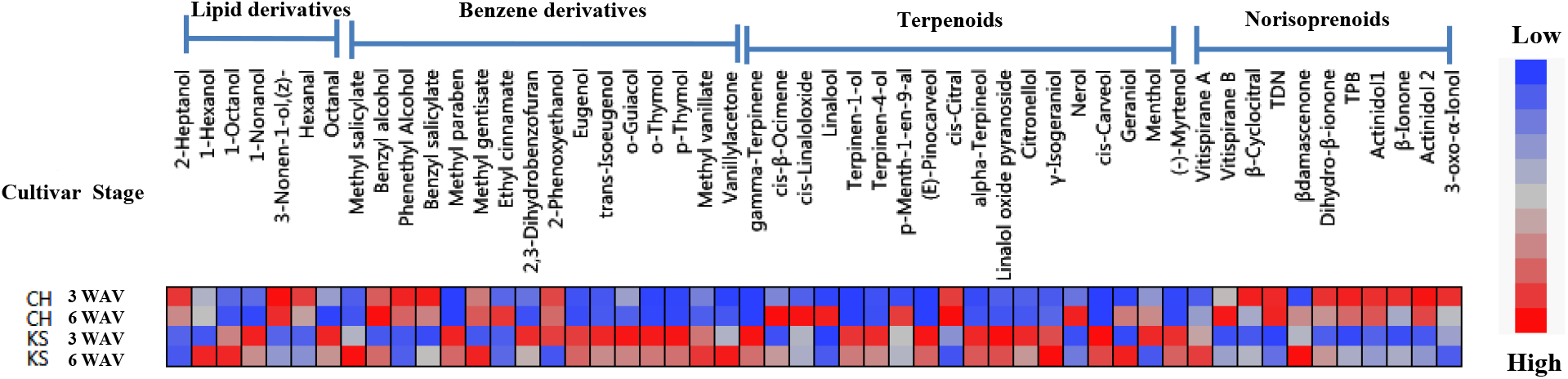
Composition of the bound volatiles in untreated Koshu (KS) and Chardonnay (CH) berries at 3 and 6 weeks after véraison [WAV]. The concentrations were determined semi-quantitatively by calculating the ratio between the area of the compound identified and the area of the internal standard 3-octanol. Data are means (n = 3). Higher concentrations for each compound are presented in red, while lower concentrations are presented in blue. There were significant differences in the concentrations of these bound volatiles between Koshu and Chardonnay (least square means Student’s t-test, *p* < 0.05).

**Table 2.** Volatile organic components (μg kg–1 of fresh weight) of Koshu and Chardonnay berries identified from the hydrolyzed fractions of precursors by gas chromatography–mass spectrometry (GC–MS).

| Chemical group/<br>Compound name | RI<sup>‡</sup> | Identification<br>assignment<sup>§</sup> | CAS number | Koshu<br>control 6 WAV | Chardonnay<br>control 6 WAV |
| --- | --- | --- | --- | --- | --- |
| **Lipid derivatives** |  |  |  |  |  |
| *C6 compounds* |  |  |  |  |  |
| 1-Hexanal<sup>†</sup> | 1098 | A | 000066-25-1 | 0.12 ± 0.11 | 0.52 ± 0.39 |
| 1-Hexanol | 1371 | A | 000111-27-3 | 43.27 ± 3.63 | 35.10 ± 5.48 |
| (E)-2-Hexen-1-ol | 1413 | A | 000928-95-0 | 19.70 ± 2.08 | 1.00 ± 0.69 |
| *Lactones* |  |  |  |  |  |
| 2-Pentylfuran | 1245 | A | 003777-69-3 | 2.19 ± 0.04 | 2.20 ± 0.06 |
| γ-Decalactone<sup>†</sup> | 2151 | A | 706-14-9 | 2.45 ± 0.33 | 3.13 ± 0.35 |
| γ-Nonalactone | 2036 | A | 000104-61-0 | ND | ND |
| *Alcohols and aldehydes* |  |  |  |  |  |
| 3-Methyl-1-butanol | 1229 | A | 000123-92-2 | 57.30 ± 1.65 | 83.37 ± 9.90 |
| Octanal<sup>†</sup> | 1301 | A | 000124-13-0 | 0.84 ± 0.32 | 0.56 ± 0.04 |
| Nonanal<sup>†</sup> | 1402 | A | 000124-19-6 | 10.11 ± 7.34 | 8.53 ± 6.28 |
| 1-Octen-3-ol | 1462 | A | 003391-86-4 | 3.77 ± 0.12 | 2.46 ± 0.36 |
| 2-Ethyl-1-hexanol | 1505 | A | 000104-76-7 | ND | ND |
| 1-Octanol<sup>†</sup> | 1570 | A | 000111-87-5 | 23.13 ± 0.93 | 15.07 ± 0.53 |
| 1-Nonanol | 1670 | A | 000143-08-8 | 3.20 ± 0.29 | 2.09 ± 0.14 |
| (Z)-3-Nonen-1-ol | 1694 | A | 010340-23-5 | 0.59 ± 0.03 | 1.01 ± 0.08 |
| *Acids and esters* |  |  |  |  |  |
| 3-Methyl-1-butanyl acetate | 1138 | A | 000123-92-2 | 2.79 ± 0.57 | 3.93 ± 0.96 |
| Hexyl acetate | 1287 | A | 000142-92-7 | 2.36 ± 0.37 | 2.51 ± 0.50 |
| 3-Hexenyl acetate | 1330 | A | 003681-71-8 | ND | ND |
| Heptyl acetate | 1383 | A | 000112-06-1 | 8.07 ± 4.09 | 15.18 ± 7.03 |
| Ethyl nonanoate<sup>†</sup> | 1542 | A | 000123-29-5 | ND | ND |
| 2-Ethylhexanoic acid <sup>*</sup> | 1956 | A | 000149-57-5 | 0.55 ± 0.83 | 0.43 ± 0.41 |
| *Miscellaneous* |  |  |  |  |  |
| Mesityl oxide<sup>†</sup> | 1155 | A | 000141-79-7 | 11.80 ± 11.28 | 6.46 ± 5.55 |
| **Benzene derivatives** |  |  |  |  |  |
| *Benzoids* |  |  |  |  |  |
| 2-Phenoxyethanol<sup>†</sup> | 2139 | A | 122-99-6 | ND | ND |
| 2,3-Dihydro-benzofuran<sup>†</sup> | >2500 | B | 496-16-2 | 42.32 ± 5.55 | 24.50 ± 3.61 |
| Ethyl cinnamate<sup>†</sup> | >2500 | B | 103-36-6 | ND | ND |
| Benzaldehyde | 1534 | A | 000100-52-7 | 0.91 ± 0.46 | 1.50 ± 0.24 |
| 4-Methyl-benzaldehyde | 1655 |  | 104-87-0 | 5.34 ± 0.26 | 4.84 ± 0.32 |
| Phenylmethyl acetate<sup>†</sup> | 1737 | C | 000140-11-4 | 7.03 ± 1.28 | 16.18 ± 3.97 |
| Methyl salicylate | 1783 | A | 000119-36-8 | 50.56 ± 17.08 | 3.53 ± 0.46 |
| 3,4-Dimethyl-benzaldehyde | 1819 | A | 005779-95-3 | 8.28 ± 0.66 | 9.07 ± 1.64 |
| 2-Phenylethyl acetate | 1823 | A | 000103-45-7 | ND | 1.16 |
| Benzyl alcohol<sup>†</sup> | 1888 | B | 000100-51-6 | 22.37 ± 8.53 | 32.28 ± 3.84 |
| Phenylethyl alcohol | 1924 | A | 000060-12-8 | 177.39 ± 23.80 | 330.55 ± 48.23 |
| Benzyl salicylate | >2500 | A | 000118-58-1 | 3.35 ± 1.07 | 4.29 ± 0.06 |
| Methyl gentisate<sup>†</sup> | >2500 | C | 002150-46-1 | ND | ND |
| Methyl paraben<sup>†</sup> | >2500 | C | 000099-76-3 | 1.12 ± 0.40 | ND |
| ***Volatile phenols*** |  |  |  |  |  |
| *p*-Cresol | 2101 | A | 000106-44-5 | 2.39 ± 0.28 | 2.14 ± 0.23 |
| Eugenol | 2177 | A | 000097-53-0 | 6.58 ± 1.31 | 1.15 ± 0.34 |
| 4-Vinylguaiacol | 2205 | A | 007786-61-0 | 269.23 ± 20.66 | 160.54 ± 17.61 |
| *p*-Thymol | 2211 | A | 003228-02-2 | 3.19 ± 0.21 | ND |
| *o*-Thymol | 2195 | A | 499-75-2 | 2.73 ± 0.42 | ND |
| *o*-Guaiacol | 1869 | A | 000090-05-1 | 5.98 ± 2.81 | ND |
| trans-Isoeugenol | 2398 | A | 005932-68-3 | 0.91 ± 0.38 | ND |
| ***Vanillin*** |  |  |  |  |  |
| Vanillyl acetone/Zingerone | >2500 | A | 000122-48-5 | 6.70 ± 1.76 | 1.72 ± 0.25 |
| Vanillin<sup>†</sup> | >2500 | A | 121-33-5 | ND | ND |
| Methyl vanillate | >2500 | A | 003943-74-6 | 33.37 ± 5.14 | 9.27 ± 2.63 |
| Acetovanillone<sup>†</sup> | >2500 | B | 498-02-2 | 3.91 ± 1.35 | 3.46 ± 0.68 |
| **Terpenoids** |  |  |  |  |  |
| (-)-Myrtenol<sup>†</sup> | >2500 | C | 19894-97-4 | 7.50 ± 0.94 | 4.52 ± 3.30 |
| Linalool oxide pyranoside | 1748 | A | 014049-11-7 | 29.11 ± 4.75 | 11.72 ± 0.81 |
| *p*-Menth-1-en-9-al<sup>†</sup> | 1619 | B | 029548-14-9 | ND | ND |
| (E)-Pinocarveol<sup>†</sup> | 1655 | C | 000547-61-5 | ND | 0.05 |
| *cis*-β-Ocimene<sup>†</sup> | 1262 | C | 003338-55-4 | ND | ND |
| gamma-Terpinene<sup>†</sup> | 1254 | B | 000099-85-4 | ND | ND |
| *cis*-Linalooloxide | 1452 | B | 1000121-97-4 | 6.38 ± 1.58 | 10.57 ± 1.35 |
| Terpinen-1-ol<sup>†</sup> | 1550 | B | 000586-82-3 | 0.52 ± 0.13 | ND |
| Linalool | 1556 | A | 000078-70-6 | 5.21 ± 1.00 | 32.98 ± 1.90 |
| Menthol<sup>†</sup> | 1617 | B | 001490-04-6 | 7.28 ± 4.42 | 3.99 ± 2.98 |
| *cis*-Citral | 1686 | A | 000106-26-3 | 2.24 ± 0.06 | 3.79 ± 0.39 |
| L-α-Terpineol | 1702 | A | 010482-56-1 | 11.96 ± 1.07 | ND |
| Terpinen-4-ol | 1607 | A | 000562-74-3 | 0.23 ± 0.03 | ND |
| γ-Isogeraniol<sup>†</sup> | 1747 | C | 013066-51-8 | 3.22 ± 0.37 | 6.10 ± 0.34 |
| Citronellol | 1777 | A | 000106-22-9 | 1.13 ± 0.10 | 0.50 ± 0.07 |
| Nerol | 1811 | A | 000106-25-2 | 8.21 ± 1.05 | 13.61 ± 1.34 |
| *cis*-Carveol | 1835 | B | 001197-06-4 | 2.94 ± 0.06 | 1.90 ± 0.01 |
| Geraniol | 1858 | A | 000106-24-1 | 23.84 ± 2.14 | 22.49 ± 1.45 |
| **Norisoprenoids** |  |  |  |  |  |
| VitispiraneA<sup>†</sup> | 1530 | B | | 12.55 ± 0.92 | 10.64 ± 2.40 |
| VitispiraneB<sup>†</sup> | 1533 | B | | 9.15 ± 0.59 | 11.75 ± 2.44 |
| β-Cyclocitral<sup>†</sup> | 1624 | B | 432-25-7 | 4.14 ± 0.25 | 3.97 ± 0.29 |
| TDN | 1748 | A | | 13.30 ± 1.48 | 27.44 ± 5.62 |
| β-Damascenone<sup>†</sup> | 1823 | A | 23726-93-4 | 30.51 ± 0.51 | 34.34 ± 4.94 |
| Dihydro-β-ionone | 1836 | A | | 0.22 ± 0.01 | 0.22 ± 0.00 |
| TPB<sup>†</sup> | 1833 | B | 1000357-25-7 | 11.95 ± 0.64 | 17.41 ± 3.06 |
| Actinidol 1<sup>†</sup> | 1935 | B | | 20.77 ± 1.06 | 31.37 ± 3.05 |
| β-ionone | 1943 | A | 79-77-6 | 4.30 ± 0.02 | 5.81 ± 1.33 |
| Actinidol 2<sup>†</sup> | 1949 | B | | 26.67 ± 0.96 | 40.79 ± 3.98 |
| 3-Oxo-α-ionol<sup>†</sup> | >2500 | B | 034318-21-3 | 1.82 ± 0.05 | 1.98 ± 0.02 |

<sup>†</sup> Where chemical standards were not available for a given compound, the total ion chromatogram area was converted using standard curves obtained for 1,1,6,-trimethyl-1,2-dihydronaphthalene (TDN) for norisoprenoids and 1-nonanol for all other compounds.
<sup>‡</sup> RI = retention index calculated in an HP-INNOWax column.
§ The reliability of the proposed identification is indicated as follows:
A- Identification carried out by comparing the gas chromatographic retention with mass spectrometric data of the pure and authentic reference compounds (pure compound available).
B- Identification tentatively carried out by comparing the gas chromatographic retention with mass spectrometric data reported in the literature (pure compound not available).
C- Pure reference compounds not available and there are no published values. Tentatively identified based on the coincidence of mass spectrometric data with the NIST Library 11.
WAV = weeks after véraison; ND = not detectable; values are means ± standard errors.

We also found higher numbers of volatile phenols, benzoids, and vanillin derivatives in Koshu berries, all of which were highly accumulated (Figure 2). For example, volatile phenols, such as eugenol, trans-isoeugenol, *o*-guaiacol, *o*-thymol, and *p*-thymol, benzoids such as methyl paraben and methyl salicylate, and vanillin derivatives, such as vanillylacetone and methyl vanillate, were more abundant in Koshu berries at the early or early and late ripening stages than in Chardonnay berries. By contrast, Chardonnay berries contained higher concentrations of benzene alcohols and esters, such as benzyl alcohol, phenethyl alcohol, and ethyl cinnamate, than Koshu berries.

#### Effects of bunch shading and ABA

We found that bunch shading significantly reduced the accumulation of norisoprenoids such as vitispirane A, TDN, TPB, and actinidols at 3 and 6 WAVs in both cultivars. However, we also observed that β-damascenone tended to increase in Koshu at 6 WAV although it was not statistically significant (Figure 3 and Supplementary Figure 1). The shading treatment also significantly affected the accumulation of terpenoids such as linalool, *y*-isogeraniol, nerol, and geraniol in both cultivars at the early and late ripening stages, and alpha-terpineol in both cultivars at 6 WAV (Figure 3). Linalool, which is the dominant terpene in Chardonnay, was most affected by the absence of light, with levels of this compound having become almost negligible at harvest. By contrast, shading had less of an effect on the other terpene compounds, with the accumulation of terpinen-1-ol, terpinenl-4-ol, and myrtenol, which were specifically found in Koshu, being slightly reduced by shading. The shading treatment also reduced the concentrations of methyl vanillate and vanillyl acetone by approximately half in Koshu, and affected the concentration of 4-vinyl guaiacol, the major volatile phenol detected, though this was not significant.

**Figure 3.**
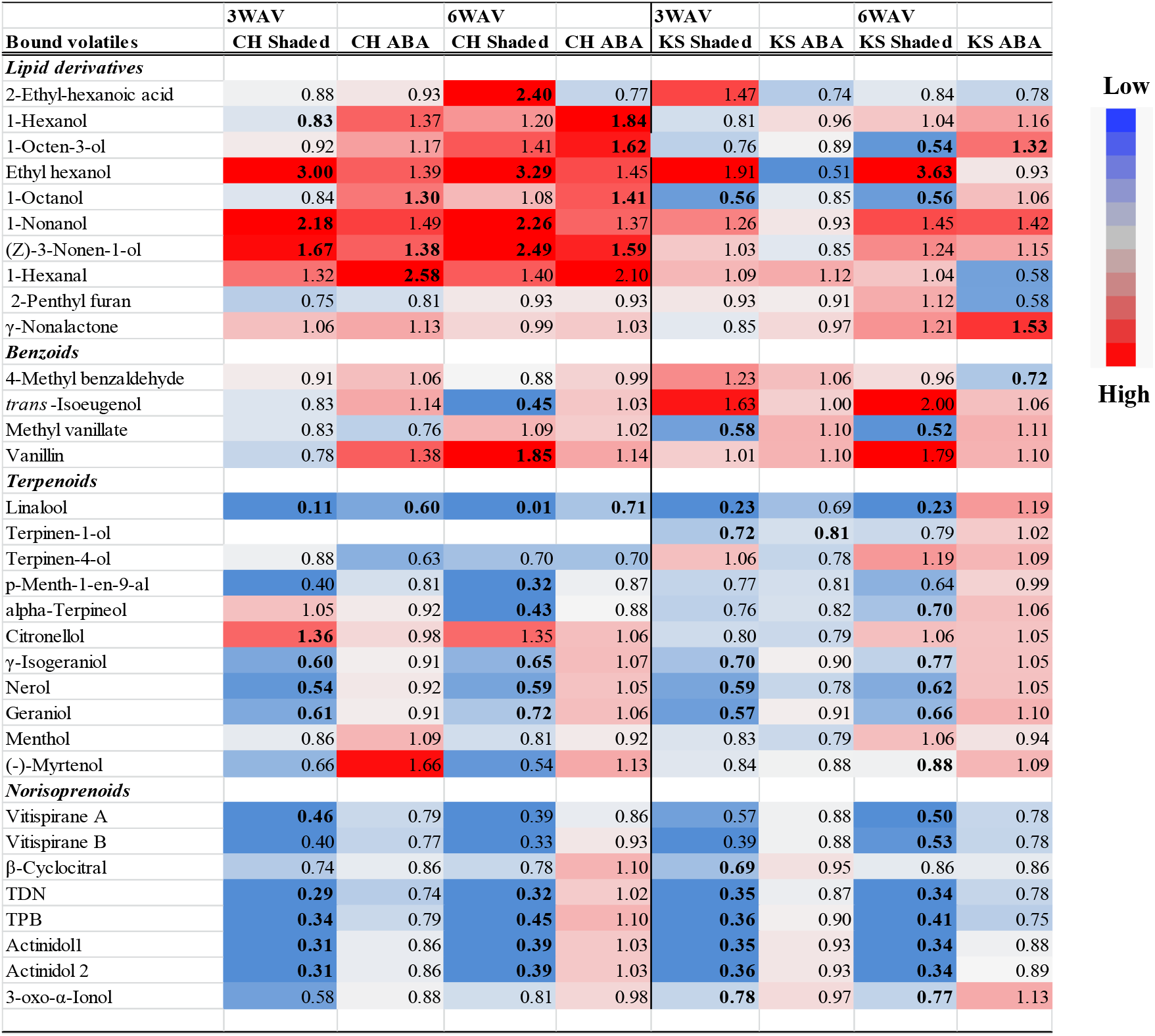
Fold-changes (treatment/control) in the concentrations of bound volatiles in Koshu (KS) and Chardonnay (CH) berries following bunch shading and abscisic acid (ABA) application. Different colors represent increases (red) and decreases (blue) in the metabolites, as indicated in the color key. Bold figures indicate that the cultivar and stage significantly increased or decreased in response to the treatment (Tukey’s honest significant difference test, *p* < 0.05). The concentrations were determined semi-quantitatively by calculating the ratio between the area of the compound identified and the area of the internal standard 3-octanol. Data are means (n = 3).

The exogenous application of ABA did not significantly affect the concentrations of terpenoids, norisoprenoids, or benzene derivatives. However, it did significantly affect the levels of lipid derivatives in Chardonnay, and the concentrations of alcohols, such as 1-hexanol, 1-octanol, 1-nonanol, 1-octen-3-ol, and 3-nonen-1-ol, which increased in both the early and late ripening stages, and hexanal, which increased at 3 WAV. In addition, ABA application tended to decrease the concentrations of all of the acetic acid esters, such as hexyl acetate, heptyl acetate, and 2-phenylethyl acetate, though this was not significant, likely due to the high levels of variation in the field. By contrast, ABA had a positive effect on the levels of *γ*-nonalactone and TPB in Koshu at the late ripening stage.

### Compositions of the free volatiles in Koshu and Chardonnay berries

The compositions of free volatiles in Koshu and Chardonnay berries differed in terms of the free forms of terpenoids (Supplementary Figure 2). Koshu berries contained higher levels of the free forms of citronellol, *γ*-isogeraniol, nerol, and geraniol, particularly at 3 WAV, none of which were detected in Chardonnay, although only *γ*-isogeraniol and geraniol were found to be significantly different between the two cultivars. By contrast, untreated Chardonnay berries contained a higher level of linalool than untreated Koshu berries. There were also significant differences between these cultivars in the composition of lipid derivatives, with Chardonnay berries containing significantly higher levels of the C_6_ compounds 1-hexanol, hexenal, 3-hexen-1-ol, acetate, and furan, 2-pentyl-, particularly at 3 WAV, and Koshu berries containing higher concentrations of ethyl hexanol, 1-octanol, and 1-nonanol, particularly at 3 WAV. These compositional differences in free form lipid derivatives were consistent with those observed for the bound forms (Figure 2). Koshu berries contained significantly higher levels of free forms of the terpenoids, *γ*-isogeraniol and geraniol, while their bound forms showed the opposite trends. Hence, the different distribution of the free and bound forms of these compounds between the cultivars were elucidated.

### Compositions of phenolic monomers and proanthocyanidins in Koshu and Chardonnay berries

#### Cultivar differences

The total phenolic compositions of the whole berries of each cultivar are listed in Supplementary Table 1. Among these, we detected some monomers (which were categorized as unknown) that we hypothesized to be putatively Koshu-or Chardonnay-specific. Koshu berries contained significantly higher levels of total phenolic acids, total hydrocinnamic acids, and total anthocyanins than Chardonnay berries (Table 3). In addition, Koshu berries tended to have higher concentrations and a different composition of proanthocyanidins than Chardonnay berries, with a significantly higher percentage of epigallocatechin (EGC) and mean degree of polymerization (mDP), and a significantly lower percentage of epicatechin-3 gallate (ECG).

**Table 3.** Concentrations (μg/g fresh weight [FW]) of total phenolic compounds extracted from the whole berries of Koshu and Chardonnay under different treatments. Berries were measured at 3 and 6 weeks after véraison (WAV) and were treated with bunch shading, the exogenous application of abscisic acid (ABA), or no treatment (control). ** Concentrations of phenolic compounds in Koshu and Chardonnay were significantly different (least square means Student’s *t*-test, *p* < 0.01). Values are means ± standard errors. Different letters indicate significant differences among treatments within each cultivar (Tukey’s honest significant difference test, *p* < 0.01).

| Phenolic monomer and Proanthocyanidin | Koshu |  |  | Chardonnay |  |  | Koshu |  |  | Chardonnay |  |  |
| --- | --- | --- | --- | --- | --- | --- | --- | --- | --- | --- | --- | --- |
|  | 3WAV | 3 WAV | 3 WAV | 3WAV | 3 WAV | 3 WAV | 6 WAV | 6 WAV | 6 WAV | 6 WAV | 6 WAV | 6 WAV |
|  | Control | Shaded | ABA | Control | Shaded | ABA | Control | Shaded | ABA | Control | Shaded | ABA |
| Total (µg/ gFW) |  |  |  |  |  |  |  |  |  |  |  |  |
| Phenolic acids ** | 27.80±2.76 | 24.01±2.87 | 30.89±3.13 | 19.10±0.86 | 18.24±0.33 | 19.72±0.76 | 29.23±2.63 | 27.12±1.06 | 27.09±1.58 | 20.53±3.35 | 20.38±5.37 | 16.68±0.72 |
| Hydrocinnamic acids ** | 242.25±30.84 | 226.00±6.73 | 193.03±11.01 | 75.16±5.41 | 78.74±3.14 | 69.75±3.46 | 199.37±39.91 | 198.75±12.18 | 168.45±13.20 | 66.52±11.13 | 69.01±22.24 | 83.08±0.56 |
| Stilbenes** | 0.06±0.06 | 0.04±0.04 | 0.11±0.06 | 0.38±0.01 | 0.18±0.01 | 0.17±0.08 | 0.07±0.07 | 0.07±0.07 | 0.04±0.04 | 0.37±0.06 | 0.20±0.17 | 0.24±0.08 |
| Flavanol | 35.45±2.75 | 23.67±2.17 | 27.63±3.02 | 30.91±1.12**A** | 6.63±0.49**B** | 26.79±1.45**A** | 38.83±2.83 | 21.84±1.08 | 38.11±3.97 | 28.86±3.91**A** | 8.85±2.19**B** | 34.00±1.06**A** |
| Anthocyanin ** | 1.47±0.21 **AB** | ND **B** | 0.96±0.05 **AB** | ND | ND | ND | 6.53±2.18 **A** | ND **B** | 6.16±0.52 **AB** | ND | ND | ND |
| Proanthocyanidin | 2098.64±723.92 | 1915.32±445.31 | 1517.34±330.80 | 1586.82±157.52 | 1306.11±60.87 | 1619.17±457.09 | 1962.98±797.77 | 1399.33±377.03 | 1366.65±199.26 | 1281.20±375.92 | 1378.70±269.42 | 1778.89±253.46 |

#### Effects of bunch shading and ABA

Neither bunch shading nor the application of ABA affected the concentrations of the major hydroxycinnamic acid derivatives, *i.e*., trans-caftaric and trans-coutaric acids. They did, however, affect some of the minor compounds (unidentified) in this class, as well as the concentrations of monomeric flavonoids. There was a significant reduction in the accumulation of flavonols under shaded conditions in Chardonnay berries and Koshu berries (Table 3), though the effect was lower on the latter cultivar, likely due to it having a different flavonol composition. Shading also reduced the concentration of stilbenes in Chardonnay berries, which contained significantly higher levels of these compounds (primarily in the form of *trans*-resveratrol) than Koshu berries.

### Principal component and linear discriminant analyses

To gain a better understanding of the differences in the aroma compounds in Koshu and Chardonnay berries at different ripening stages, and the effects of shading and ABA application on these, we subjected all of the variables (metabolites in this case) of the bound volatiles to PCA and LDA (Figure 4). Principal component 1 (PC1) contributed 31.5% and principal component 2 (PC2) contributed 18.5% of the total variance. A scatter plot of 35 samples according to PC1 and PC2 showed a clear separation of the volatile profiles between the two cultivars along PC1 (Figure 4A). The separation by the treatments was less clear, but we were able to observe some differences along PC2, particularly between shaded berries and all other treatments. PC1 was highly correlated with several minor terpenoids and benzene derivatives, and was considered to characterize the bound volatiles in Koshu berries (Figure 4B). Among these, (E)-pinocarveol, cis-carveol, *y*-terpinene, terpinen 1-ol, and methyl paraben, some of which were tentatively identified in this study, were found to be Koshu-specific. We also constructed an LDA model for 13 compounds selected by forward stepwise regression (Figure 4C). This model successfully classified berries that had been subjected to different treatments (accuracy of classification: 100%; prediction score: 88.6%), with shaded samples having higher prediction scores (100%). Several terpenoids and norisoprenoids, such as actinidol 1, terpinen-1-ol, and geraniol, had high positive CV1 scores (Supplementary Table 2). ABA-treated samples were also discriminated by the LDA model (Figure 4C), with the selected volatile compounds, such as hexanal, nonanol, and y-nonalactone, contributing to the discrimination (Supplementary Table 2).

**Figure 4.**
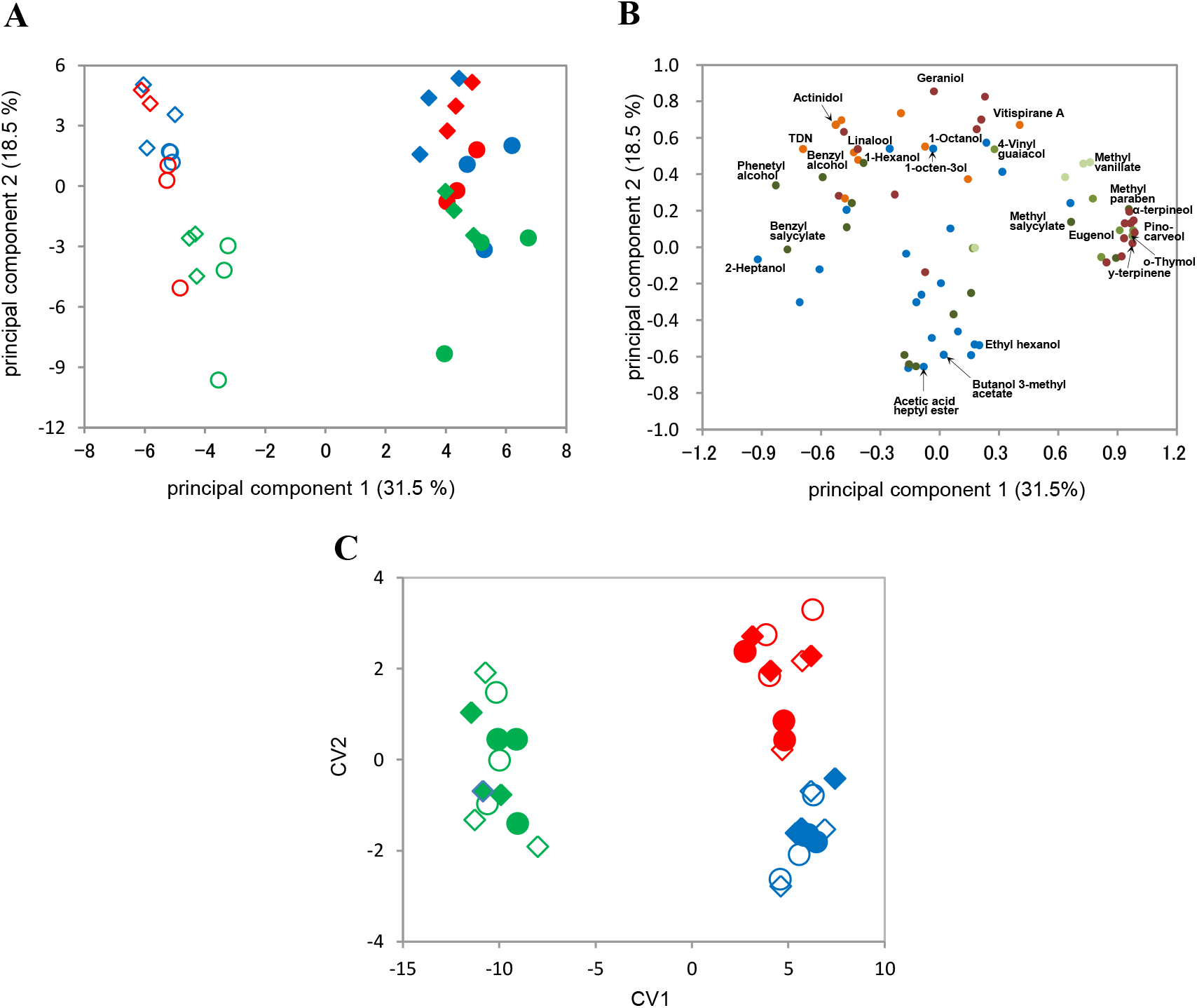
Principal component analysis (PCA) and linear discriminant analysis (LDA) of the bound volatiles in the berries of Koshu and Chardonnay under different treatments. (A) The first two component scores of the samples for PCA. (B) Loading plot of the variables, showing the positions of the compounds classified as lipid derivatives (blue), terpenoids (purple), benzoids (dark moss green), volatile phenols (moss green), vanillins (pale moss green), and norisoprenoids (orange); selected variables for interpreting the score plot are labeled. (C) Scatterplot of the best two discriminant functions in the LDA model, which included the concentrations of 13 selected bound volatiles in the berries discriminated by cultivar (Chardonnay, empty marker; Koshu, filled marker) and treatment (control, blue; shade, green; ABA application, red). Samples that were collected at 3 and 6 weeks after véraison (WAV) are shown as circles and diamonds, respectively.

## DISCUSSION

In this study, we comprehensively analyzed and compared the secondary metabolite profiles of Koshu and Chardonnay wine grapes, with a particular focus on volatile compounds (both glycosidically bound and free) and phenolic compounds. We also investigated how the aroma and flavor characteristics of these cultivars are influenced by bunch shading and the exogenous application of ABA at different ripening stages.

### Cultivar differences

When investigating the aroma compounds in the berries of these cultivars, we deliberately focused on the bound volatiles, as these are the main contributory factors to the quality of wine. We found that there were remarkable differences in the composition of bound volatiles between the two cultivars, with Koshu berries containing significantly more volatile phenols and vanillins, and Chardonnay berries containing significantly more norisoprenoids and benzoids (Figure 1). Consistent with the result, Sefton *et al*.^9^ observed that Chardonnay juices contain remarkably high ratios of norisoprenoids in the bound form, suggesting that these make a significant contribution to the sensory properties of their wine. On the other hand, eugenol, vanillin, and guaiacol in volatile phenols are released from the bound forms contained in the ripening berries after the vinification processes and aging, and contribute to spicy, roasted, and oak wood aromas.^38^ Therefore, the remarkably high concentrations of these precursors in Koshu berries may contribute to the sensory properties of Koshu wine, giving it these aroma attributes even if it is not aged in barrels. Interestingly, Koshu berries also contained a large number of minor terpenoids that have rarely been found, particularly in *V. vinifera* cultivars, suggesting the unique characteristics of the bound terpenoid profile of Koshu.

In the PCA, there was a clear separation of the volatile profiles between the two cultivars along PC1, which was highly correlated with several minor terpenoids and benzene derivatives that were found to be Koshu-specific (Figure 4 and Supplementary Figure 1). Eugenol and isoeugenol, which are abundant volatile phenols in Koshu berries, are also produced by plants as defense compounds against animals and microorganisms. In addition, Koshu berries contained remarkably high concentrations of methyl salicylate, which is the central hormone that systemically induces plant resistance to biotrophs^39^. We therefore hypothesize that these volatile phenols and benzene derivatives may be necessary for defense against diseases, making Koshu more resistant to diseases than pure-bred *V. vinifera* cultivars such as Chardonnay. This unique profile of Koshu that is distinct from that of *V. vinifera* also infers that these traits have been inherited from its East Asian wild species parent.

### Effects of bunch shading and ABA

The LDA model of aroma potential in grape berries was able to discriminate the quality of berries grown under different regimes, with shaded samples having the highest prediction scores (100%), indicating the remarkable influence of light on the aroma potential of berries (Figure 4C). In both cultivars, bunch shading significantly decreased TSS concentrations and significantly increased TA levels, retarding the ripening process, as reported previously^40^. Shaded berries also generally contained lower concentrations of norisoprenoids and terpenoids (Figure 3), suggesting that shading has a negative effect on their quality. In the vineyard, reduced sunlight exposure appears to reduce the production of carotenoids in the berries, which subsequently decreases the levels of norisoprenoids in the finished wine. However, the most abundant norisoprenoid in Koshu, β-damascenone, tended to increase under shading, and that β-damascenone is an important component of wine and is known to have significant indirect impacts on wine aroma and quality,^16^ although its direct effects are still being investigated. Rain-cut shade is used in Japanese vineyards to prevent the spread of disease caused by rain and certain types of shading have been reported to reduce the phenolic contents in Koshu, improving the quality of the berries^41^. We did not observe a significant reduction in the total contents of phenolic compounds except flavonol and anthocyanins as a result of shading in this study. However, we did find that some of volatile derivatives from phenols, such as methyl vanillate, vanillyl acetone and 4-vinyl guaiacol decreased, particularly in Koshu berries. 4-vinyl guaiacol is responsible for smoky and phenolic notes, and is considered to be an off-flavor when present at a high abundance in wine. Thus it was suggested that shading may also have positive effects for mitigating some of the unfavorable flavors that are caused by volatile phenol derivatives in this cultivar. The LDA model could also discriminate ABA-treated samples (Figure 4C). The exogenous application of ABA caused a yellowing of the skins of Chardonnay berries, suggesting accelerated ripening, but had no significant effect on the other maturation indices, supporting the previous findings of Koyama *et al*.^19^ ABA application had no significant effect on the concentrations of terpenoids, norisoprenoids, or benzene derivatives. However, it did have significant effects on the levels of lipid derivatives and the concentrations of alcohols, with hexanal, nonanol, and *y*-nonalactone contributing to the discrimination by the LDA model. This supports the previous finding that ABA application improves the accumulation of C6 aroma compounds derived from fatty acids.^20^ Furthermore, Kalua *et al*.^42^ reported that free C_6_ volatile compounds change from acetate esters to aldehydes and finally to alcohols, with a predominance of alcohols in the late stages of berry development in Cabernet Sauvignon. The pattern of lipid derivative accumulation that was observed in the present study suggests that ABA application accelerates ripening, particularly in Chardonnay. Moreover, the observed differences in the volatile profiles of the two cultivars suggest that ABA affects fatty acid metabolism (the lipoxygenase [LOX] pathway). Ethylene signaling has been reported to be involved in the regulation of the flavor pathway during the late ripening stage of grape berries,^3^ and an interaction between ethylene signaling and ABA application has been reported previously.^19^ Therefore, further research is required to elucidate the relationship between environmental cues and hormone levels in the berries, particularly during the late ripening stage.

## CONCLUSION

From our analyses, we were able to discriminate the flavor precursor profiles of white wine grapes by cultivar and the growth environment. Koshu was found to possess a unique profile of quality-related metabolites in its berries, as well as a high abundance of minor terpenoids and volatile phenolic compounds, suggesting their role in stress adaptation to the Japanese climate. Furthermore, we found that Koshu and Chardonnay exhibited different responses to bunch shading, which were related to compositional differences in the bound flavor compounds they contain. The unique secondary metabolite profile and responsiveness to the environment in Koshu could be important not only for developing various styles of Koshu wines with interesting aromas and flavors, but also for producing new cultivars with improved quality and cultural characteristics.

## Acknowledgment

This work was supported by a Grant-in-Aid for Scientific Research (grant no. 15K07306) from the Japan Society for the Promotion of Science.

